# The effects of genome size on cell size and the functional composition and morphology of leaves: a case study in *Rhododendron* (Ericaceae)

**DOI:** 10.1101/2023.10.13.562260

**Authors:** Arezoo Dastpak, Monica Williams, Sally Perkins, John A. Perkins, Charles Horn, Patrick Thompson, Connor Ryan, Juliana Medeiros, Yi-Dong An, Guo-Feng Jiang, Kevin A. Simonin, Adam B. Roddy

## Abstract

**Background and Aims:** Despite the predominance of scaling photosynthetic metabolism by two-dimensional leaf surface area, leaves are three-dimensional structures composed of multiple tissues that directly and indirectly influence photosynthetic metabolism. The structure of leaf surfaces for CO_2_ diffusion and light transmission and the internal volume of tissues that process energy and matter work together to control rates of resource acquisition and turnover. Here we investigate the influence of cell size and packing density on resource acquisition as measured by surface conductance to CO_2_ and water vapor and on resource turnover as measured by leaf water turnover time.

**Methods:** We sampled wild and cultivated congeneric species in the genus *Rhododendron* (Ericaceae) and measured genome size, anatomical traits related to cell sizes and packing densities, and morphological traits related to water content and dry mass allocation.

**Results:** Among *Rhododendron*, anatomical traits related to cell size and morphological traits related to water content and dry mass investment varied largely orthogonally to each other, allowing for many combinations of leaf traits to exist. However, there was a strong, negative relationship between the leaf water residence time (τ) and the maximum leaf surface conductance per leaf volume (*g*_*max,vol*_), both of which are influenced by cell size and cell packing densities.

**Conclusions:** Despite leaf function being controlled by many potential combinations of leaf cell- and tissue-level traits, cell size has a pervasive effect on leaf function. Small cells allow for higher diffusion of CO_2_ and water vapor per unit leaf volume (*g*_*max,vol*_) even at constant leaf thickness, but small cells also result in shorter leaf water residence times (τ). The strong tradeoff between *g*_*max,vol*_ and (τ) illuminates how genome size-cell size allometry influences the fast-slow continuum of plant carbon and water economy.

## Introduction

Because leaves are responsible for regulating the exchange of energy and matter between plants and the atmosphere, they play a central role in whole-plant strategies for resource acquisition and growth, thereby impacting ecosystem function and climate (Boyce and Lee 2010; Bonan *et al*. 2014; Franks *et al*. 2017). Quantifying structural constraints on leaf function has, therefore, been a central goal in plant functional ecology (Bazzaz *et al*. 1987; Chapin 1989; Wright *et al*. 2004; Violle *et al*. 2007; Reich 2014). One example, the leaf economics spectrum (LES), identifies a suite of coordinated traits describing tradeoffs in resource allocation between photosynthesis and biomechanical structure that impact leaf longevity and whole-plant relative growth rate (Reich *et al*. 1992). This resource allocation tradeoff is typified by a single trait, specific leaf area (*SLA*) or its reciprocal, leaf dry mass per unit projected surface area (*LMA*) (Wright *et al*. 2004). Although *SLA* and *LMA* are widely used to describe leaf-level metabolism, one aspect of the LES framework that often complicates interpretations is the difference observed when indices of leaf metabolism (e.g. photosynthetic rate, respiration rate, stomatal conductance) are normalized by leaf dry mass versus by the surface area for light interception, i.e. one-sided projected leaf surface area.

Within leaves, cells are organized into tissues that must efficiently fill the leaf volume and accomplish multiple functions, including scattering and absorbing light, facilitating CO_2_ diffusion into the mesophyll cells for photosynthesis, and remaining biomechanically robust (Smith *et al*. 1997; Roderick, Berry, Saunders, *et al*. 1999; Théroux-Rancourt *et al*. 2021; Borsuk *et al*. 2022; Treado *et al*. 2022). None of these functions are performed solely by individual cells: leaf performance depends on the whole leaf, which is composed of multiple tissues and their constituent cells. This hierarchical organization allows for variation at each level of organization to modify the effects of variation at lower levels on whole organ function. Because filling a tissue volume requires allocation of resources, such as carbon and water, to cells that occupy space (Roderick, Berry, Noble, *et al*. 1999), leaf function depends not only on the two-dimensional surfaces across which energy and matter are exchanged but also on the three-dimensional tissue volumes–and their component masses–where energy and matter are stored and metabolized (Roderick, Berry, Saunders, *et al*. 1999; Théroux-Rancourt *et al*. 2021, 2023; Trueba *et al*. 2022).

Though fluxes of CO_2_ and water between plants and the atmosphere are typically calculated per unit leaf surface area, the turnover times of matter that ultimately scale with leaf longevity and whole-plant relative growth rate depend on the leaf volume that is supplied by a given surface area (Roderick, Berry, Noble, *et al*. 1999; Roderick, Berry, Saunders, *et al*. 1999; Shipley *et al*. 2006). In order for CO_2_ to enter the mesophyll cells it must cross the leaf surface and diffuse through the intercellular airspace volume inside the mesophyll tissue, both of which are constructed to optimize the supply of CO_2_ to sites of photosynthetic metabolism (Théroux-Rancourt *et al*. 2021). Stomata in the leaf epidermis control the surface conductance for net CO_2_ transport into the intercellular airspaces of the leaf, and the size and spatial arrangement of cell surfaces that define the intercellular airspace volume supplied by stomata determine the total surface area available for CO_2_ uptake (Trueba *et al*. 2022; Théroux-Rancourt *et al*. 2023). This suggests that scaling from cell- and tissue-level traits to whole-leaf traits should account for variation in the total surface area available for CO_2_ uptake per unit leaf volume (Earles *et al*. 2018; Théroux-Rancourt *et al*. 2021; Borsuk *et al*. 2022). The development of the LES has incorporated total leaf volume insofar as leaf mass per area increases with leaf thickness (Shipley 1995; Poorter *et al*. 2009; De La Riva *et al*. 2016). However, leaf thickness is only one component of *LMA* and may account for a small fraction of interspecific variation in *LMA* (Witkowski and Lamont 1991; Shipley 1995; De La Riva *et al*. 2016; John *et al*. 2017). As a result, the total mesophyll surface area per unit leaf volume can vary independently of LMA.

These simple biophysical effects of surface area and volume have a strong influence on leaf structure and function. Thinner leaves that have a higher ratio of projected surface area to volume (*SA*_*leaf*_*/V*_*leaf*_) can exchange more energy and matter with the atmosphere per unit volume than thicker leaves can. Similarly, a given leaf volume can maintain higher CO_2_ diffusion if its constituent cells are smaller because smaller cells allow for a higher mesophyll surface area per unit leaf volume (John *et al*. 2013; Théroux-Rancourt *et al*. 2021; Borsuk *et al*. 2022).

Importantly, the minimum possible cell size, the maximum possible cell packing density, and mesophyll surface area for CO_2_ diffusion is fundamentally limited by the size of the genome (Cavalier-Smith 1978; Beaulieu *et al*. 2008; Šímová and Herben 2012; Simonin and Roddy 2018; Roddy *et al*. 2020; Théroux-Rancourt *et al*. 2021; Borsuk *et al*. 2022; Jiang *et al*. 2023). Mechanistically, genome size defines the minimum limit of cell volume and the maximum limit of cell packing densities, but beyond these bounds, cell sizes and packing densities can vary dramatically depending on leaf morphology, development, or abiotic conditions (Carins Murphy *et al*. 2016; Roddy *et al*. 2020; Jiang *et al*. 2023). Thus, scaling from individual cells to whole leaves allows for many possible combinations of leaf architectures. For example, both thick leaves and thin leaves could be composed of either small or large cells, but thin leaves and/or leaves with small cells would be capable of higher rates of energy and matter exchange than thick leaves and/or leaves with large cells.

Because metabolic processes occur in an aqueous environment, photosynthetic capacity is affected not only by the rates of CO_2_ diffusion into the leaf but also by the leaf water content (Roderick, Berry, Noble, *et al*. 1999; Roderick, Berry, Saunders, *et al*. 1999). Recent incorporation of water mass into the LES has shown that it is a strong predictor of maximum photosynthesis (Huang *et al*. 2020; Wang *et al*. 2022). Because the vacuole comprises the majority of the cell volume, larger cells have absolutely higher potential metabolic capacity than smaller cells, and leaves with more water content have higher metabolic capacity than leaves with lower water content. All else being equal, a thick leaf would have higher water content per leaf surface area than a thin leaf and, by extension, a higher potential photosynthetic capacity per unit projected leaf surface area. However, a thick leaf would typically have lower potential photosynthetic capacity per leaf volume. Furthermore, at constant leaf surface conductance, increasing leaf thickness would increase leaf water content per projected leaf area but result in slower turnover of leaf water. The turnover time (τ, also called the leaf water residence time) is defined as the ratio of water loss to water content (see Methods) and defines how rapidly the leaf water pool is replaced (Farquhar and Cernusak 2005; Simonin *et al*. 2013; Roddy *et al*. 2023). Because high water content buffers physiological responses to changes in the environment, lengthy τ is indicative of non-steady state physiology that responds minimally to environmental fluctuations (Hunt and Nobel 1987). As such, τ may be a powerful metric for characterizing ecological strategies, such as resource conservatism or resource acquisitiveness along the fast-slow continuum of plant strategies (Reich 2014).

Here we test how genome size-cell size allometry influences leaf functional anatomy using a group of closely related *Rhododendron* species. The genus *Rhododendron* is often divided into four groups: evergreen azalea, elepidote, lepidote, and deciduous azaleas, and these groups have been shown to vary in ploidy and, thus, genome size (Schepper *et al*. 2001; Jones *et al*. 2007; De *et al*. 2010; Hu *et al*. 2023). Using this variation in genome sizes among *Rhododendron*, we predicted (1) that mature cell sizes and packing densities would be correlated with genome size but would always be larger than the minimum cell size defined by the size of meristematic cels, (2) that morphological traits (lamina thickness, LMA, leaf water content) would be correlated with each other as predicted by the LES but largely independent of variation in anatomical traits (e.g. cell sizes and packing densities), but (3) that cell size would nonetheless mediate the effects of leaf morphological traits on maximum metabolic capacity. Testing these hypotheses helps clarify the various ways that cells, tissues, and whole leaves can be built and the potential effects of variation at different levels of organization on leaf ecological strategies.

## Methods

### Rhododendron diversity and plant material

In total, we measured 162 samples from 147 *Rhododendron* accessions (accessioned material in botanical gardens or wild-growing plants). For 15 of the naturally occurring azaleas, we sampled the same individual plants in years 2022 and 2023, in order to validate our measurements of genome size from 2022. These samples include 65 deciduous azaleas, of which 42 were from identified species growing in the field, 17 were naturally occurring hybrids whose parentage was based on intermediate phenotypes, and six were from artificially generated hand crosses (Table S1). In addition to naturally occurring plants, we also collected artificially created hybrids, as well as a variety of *Rhododendron* taxa from botanical gardens throughout the U.S., including the University of California Botanic Garden in Berkeley, CA, the Rhododendron Species Foundation Garden in Federal Way, WA, the Holden Arboretum in Kirtland, OH, the New York Botanic Garden in New York, NY, and the Davis Arboretum of Auburn University. In total, sampled *Rhododendron* included representatives from four of the currently recognized *Rhododendron* subgenera: Tsutrusi, Rhododendron (including tropical Vireya), Pentanthera, and Hymenanthes (Xia *et al*. 2022). However, not all traits were measured on all samples; in particular, tropical Vireya were not measured for morphological traits, such as leaf thickness, water content, and *LMA*.

### Genome size

To determine genome size by flow cytometry, we followed standard protocols for measuring genome size in plants (Dolezel *et al*. 2007; Pellicer and Leitch 2014). Approximately 50–100 mg of young and fresh leaf tissue was finely chopped over ice using a fresh razor blade along with fresh standard leaf material [*Zea mays* L., 1C = 2.71 pg; Lysak and Dolezel (1998)] in 2000 μl ice-cold Galbraith’s buffer (45 mM MgCl_2_, 20 mM MOPS, 30 mM sodium citrate, 0.1% (v/v) Triton X-100, pH 7.0, (Galbraith *et al*. 1983)). Seeds of the plant standard were generously provided by the Institute of Experimental Botany, Czech Academy of Sciences. After filtering the homogenate through a 30-μm nylon mesh filter (CellTrics™, Sysmex, Germany), 50–100 μg/mL propidium iodide was added. Samples were incubated on ice for 15 minutes prior to analysis. Flow cytometry was performed using a BD Accuri C6 Flow Cytometer (BD Biosciences, San Jose). For each unknown sample, at least 5000 nuclei were counted, with a coefficient of variation <5% for measured peaks. The 2C-value representing the genome size was determined as (Pellicer and Leitch 2014):

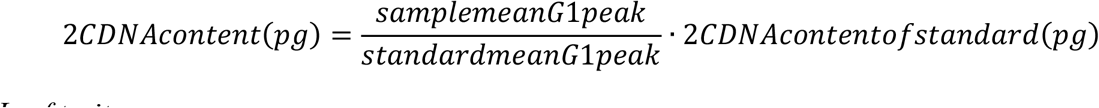

### Leaf traits

Freshly cut shoots were kept in humid plastic bags during transport back to the laboratory prior to sample processing. For each plant, we selected two leaves for analysis, both of which were measured for leaf thickness (*T*_*l*_) in three locations using a digital thickness gauge (resolution: 0.01 mm; Mitutoyu 700-118-20), taking care to avoid prominent veins. One of these leaves was weighed for fresh mass, scanned for leaf area and shape, and dried for at least 72 hrs at 70°C for subsequent dry mass measurement (resolution: 0.001 g; Sartorius, Göttingen, Germany). The other leaf was used for anatomical measurements. Approximately 1 cm^2^ sections of leaves were cleared by incubating them at 70°C in a 1:1 solution of H_2_O_2_ (30% hydrogen peroxide) and CH_3_COOH (100% acetic acid) for 24 hrs. The sections were then thoroughly rinsed in water, and their epidermises carefully separated from the layer of mesophyll and veins using a paintbrush. The epidermal layers were then stained with Safranin O (1% w/v in water) for 5-10 minutes, followed by a wash with water and a subsequent staining with Alcian Blue (1% w/v in 50% v/v ethanol) for 1 min and a rinse in 85% ethanol. The stained layers were then mounted onto microscope slides using CytoSeal (Fisher Scientific). Images were captured at varying magnifications (10x, 20x, or 40x) using a compound microscope equipped with a digital camera (Raspberry Pi High Quality Camera, Raspberry Pi Foundation). Both the abaxial and adaxial surfaces of the leaves were imaged for all species.

We used ImageJ (Rueden *et al*. 2017) to measure leaf anatomical traits. Guard cell length (*l*_*g*_) was measured on at least 10 stomata per leaf from images taken at 40x magnification. The two-dimensional areas of epidermal pavement cells (*A*_*ec*_) and stomatal guard cells (*A*_*s*_) were measured by tracing the outlines of at least ten pavement cells or stomatal complexes (two guard cells) for each sample on 40x images. Stomatal (*D*_*s*_) and epidermal pavement (*D*_*ec*_) cell densities were measured on 20x or 40x images by counting all the cells of each cell type in a field of view and dividing by the area of the field of view. For most samples, the field of view was the entire image, but when the entire image was not in focus, only the image area in focus was used. We measured leaf vein density (*D*_*v*_) as the total length of veins in an image divided by the dimensions of the image. To compare our anatomical measurements with previously published data for angiosperms, we used the dataset of *l*_*g*_, *D*_*s*_, and *D*_*v*_, compiled by Jiang et al. (2023), which included data for 836 species from 126 families with 289 species that had both *l*_*g*_ and *D*_*s*_ measurements. Meristematic cell volumes as a function of genome size were taken from Šímová and Herben (2012). Using these measured volumes of meristematic cells, we approximated the maximum two-dimensional cross-sectional area of a spherical meristematic cell by calculating the cross-sectional area of a sphere with the same volume. We also estimated the maximum packing density of spherical meristematic cells as the reciprocal of the cross-sectional area of a spherical meristematic cell [Théroux-Rancourt *et al*. (2021).

### Data transformations and analyses

From measurements of leaf fresh mass (*M*_*f*_), dry mass (*M*_*d*_), and leaf area (*A*_*L*_), we calculated the leaf mass per area (*LMA*) as:

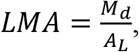

and leaf water content per unit area (*W*_*area*_) was calculated as:

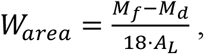

where 18 represents the conversion from grams of water to moles of water. Leaf water content per unit dry mass (*W*_*mass*_) was calculated as:

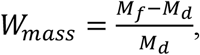

and the proportion of fresh leaf mass that is water (*W*_*prop*_) was calculated as:

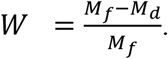

Maximum stomatal conductance to water vapor was calculated from measurements of *l*_*g*_ and *D*_*s*_ according to Franks and Beerling (2009):

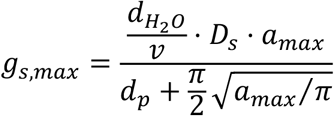

where *d*_*H2O*_ is the diffusivity of water vapor in air (0.0000249 m^2^ s^-1^), *v* is the molar volume of air normalized to 25 ^∘^C (0.0224 m^3^ mol^-1^), *d*_*p*_ is the depth of the stomatal pore, and *a*_*max*_ is the maximum stomatal pore area. The depth of the stomatal pore, *d*_*p*_, was assumed to be equal to the width of one guard cell, which was taken as 0.36 ⋅ *l*_g_ (de Boer, Price, *et al*. 2016). The maximum area of the open stomatal pore, *a*_*max*_, was approximated as *π*(*p*/2)^2^ where *p* is stomatal pore length and was approximated as *l*_g_/2.

Maximum stomatal conductance, *g*_*s,max*_, was used to calculate the leaf water residence time as:

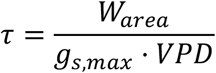

with *VPD* = 1 kPa (Roddy *et al*. 2023). Because stomatal conductance under natural conditions is likely never near its anatomically defined maximum of *g*_*s,max*_, this estimate of τ is likely much shorter than any τ encountered in nature. Nonetheless, it can be used to compare how *W*_*area*_ and leaf anatomy influence the temporal dynamics and responsiveness of leaf physiology to variation in the environment.

To express these variables on a volume basis, we calculated the ratio of leaf surface area to volume as:

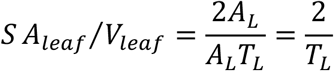

where the 2 accounts for the total leaf surface area of a planar leaf being twice the measured projected surface area (*A*_*L*_) and *T*_*L*_ is leaf thickness. *SA*_*leaf*_*/V*_*leaf*_ simplifies to being a scalar function of only leaf thickness, as shown. While both surfaces of the leaf are used for energy exchange, in hypostomatous leaves only one surface is used for gas exchange. To express *g*_*s,max*_ on a volumetric basis accounting for the fact that gas exchange occurs on only one surface, we calculated *g*_*max,vol*_ as:

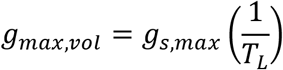

where *T*_*L*_^-1^ is equivalent to half of *SA*_*leaf*_*/V*_*leaf*_ to account for gas exchange occurring on only one leaf surface.

To determine how *SA*_*leaf*_*/V*_*leaf*_ and cell size interact to influence *g*_*max,vol*_ and τ, we examined how traits covaried with regression residuals. We first used standard major axis regression (Warton *et al*. 2012) to determine the relationship between *g*_*max,vol*_ or τ and *SA*_*leaf*_*/V*_*leaf*_. The residuals of these regressions signify the variation in *g*_*max,vol*_ or τ that is unexplained by *SA*_*leaf*_*/V*_*leaf*_. We then tested whether these residuals were related to cell size. To minimize autocorrelation due to guard cell size being used in the calculation of *g*_*s,max*_, we used instead epidermal cell size, which nonetheless scaled with *l*_*g*_ (*R*^*2*^ = 0.32, *P* < 0.0001). There were significant negative relationships between epidermal cell size and the residuals of *g*_*max,vol*_ or τ against *SA*_*leaf*_*/V*_*leaf*_ (*g*_*max,vol*_: *R*^*2*^ = 0.11, *P* < 0.0001; τ: *R*^*2*^ = 0.08, *P* = 0.003), such that leaves with smaller cells tended to have positive residuals in the *g*_*max,vol*_ relationship (i.e. above the regression line in Figure 5a) and negative residuals in the τ relationship (i.e. below the regression line in Figure 5c). To show the effects of cell size on the *g*_*max,vol*_ or τ versus *SA*_*leaf*_*/V*_*leaf*_ relationships, we calculated epidermal cell size isoclines for four epidermal cell sizes (400, 800, 1200, 1600 μm^2^) by using the linear regression between the residuals and epidermal cell size; the residual values for each cell size was then added to the regression relationship of *g*_*max,vol*_ or τ versus *SA*_*leaf*_*/V*_*leaf*_ to generate the isoclines, which represent the average effect of cell size on the *g*_*max,vol*_ or τ versus *SA*_*leaf*_*/V*_*leaf*_ relationship (Figure 5b,d).

All statistical analyses were conducted in R (v. 4.1.2) (R Core Team 2018). Because the plants we sampled included putative interspecifc interploidy hybrids, experimentally produced crosses with unclear taxonomic identity, and individuals sampled across species’ ranges, we aggregated data to the individual plant level without reference to taxonomic rank (i.e. we did not calculate species means). Traits were log-transformed prior to most analyses when doing so improved normality. We used standard major axis (SMA) regression (R package ‘smatr’) to determine the scaling relationships between traits (Warton *et al*. 2012) and show confidence intervals around SMA regressions by bootstrapping the SMA regressions 1000 times. We used slope tests, implemented in ‘smatr’, to compare slopes, and we report *P*-values for whether the slopes are significantly different or not. Principal components analysis (PCA) using the R function ‘princomp()’ was used to determine multivariate trait covariation. Traits were log-transformed, centered, and scaled prior to calculating principal components. To partition variance explained among multiple factors, we used the function ‘varpart()’ in the R package ‘vegan’ (Oksanen *et al*. 2007).

## Results

### Genome size diversity among Rhododendron

Consistent with previous studies of *Rhododendron* ploidy and genome size, our sampling found a range of 2C genome sizes among *Rhododendron*, varying from 1.10 pg for *R. obtusum* to 3.90 pg for *R. leucogigas* (Table S1). These fell into three distinct groups that displayed 2C genome size values consistent with differences in ploidy (Figure S1). Notably, some of the co-occurring deciduous azaleas included differences in genome size that would be consistent with differences in ploidy (Table S1) [**Supplementary Information**].

### Scaling of genome size with cell sizes and packing densities

Among *Rhododendron* and angiosperms more broadly, the volumes and two-dimensional sizes of mature stomatal guard cells and epidermal pavement cells were always larger than the volumes and two-dimensional sizes of meristematic cells, and the packing densities of guard cells and epidermal cells were always lower than the packing densities of meristematic cells (Figure 1). Nonetheless, genome size was a strong predictor of guard cell volumes among Rhododendron (*R*_*2*_ = 0.20, *P* < 0.0001) and angiosperms more broadly (*R*_*2*_ = 0.38, *P* < 0.0001; Figure 1a). The two-dimensional sizes of stomatal guard cells (*R*_*2*_ = 0.20, *P* < 0.0001) and epidermal pavement cells (*R*_*2*_ = 0.18, *P* < 0.0001) also scaled with genome size among *Rhododendron*, and stomatal guard cells were general smaller than epidermal pavement cells (Figure 1b). Guard cell size also scaled strongly with genome size among a broader set of angiosperms (*R*_*2*_ = 0.38, *P* < 0.0001; Figure 1b). Though among angiosperms stomatal density was scaled negatively with genome size (*R*_*2*_ = 0.23, *P* < 0.0001), among *Rhododendron* there was no significant relationship between stomatal density and genome size (*P* = 0.1; Figure 1c). However, epidermal cell packing density, which was higher than stomatal density, scaled negatively with genome size among genus *Rhododendron* (*R*_*2*_ = 0.10, *P* < 0.04; Figure 1c).

**Figure 1.**
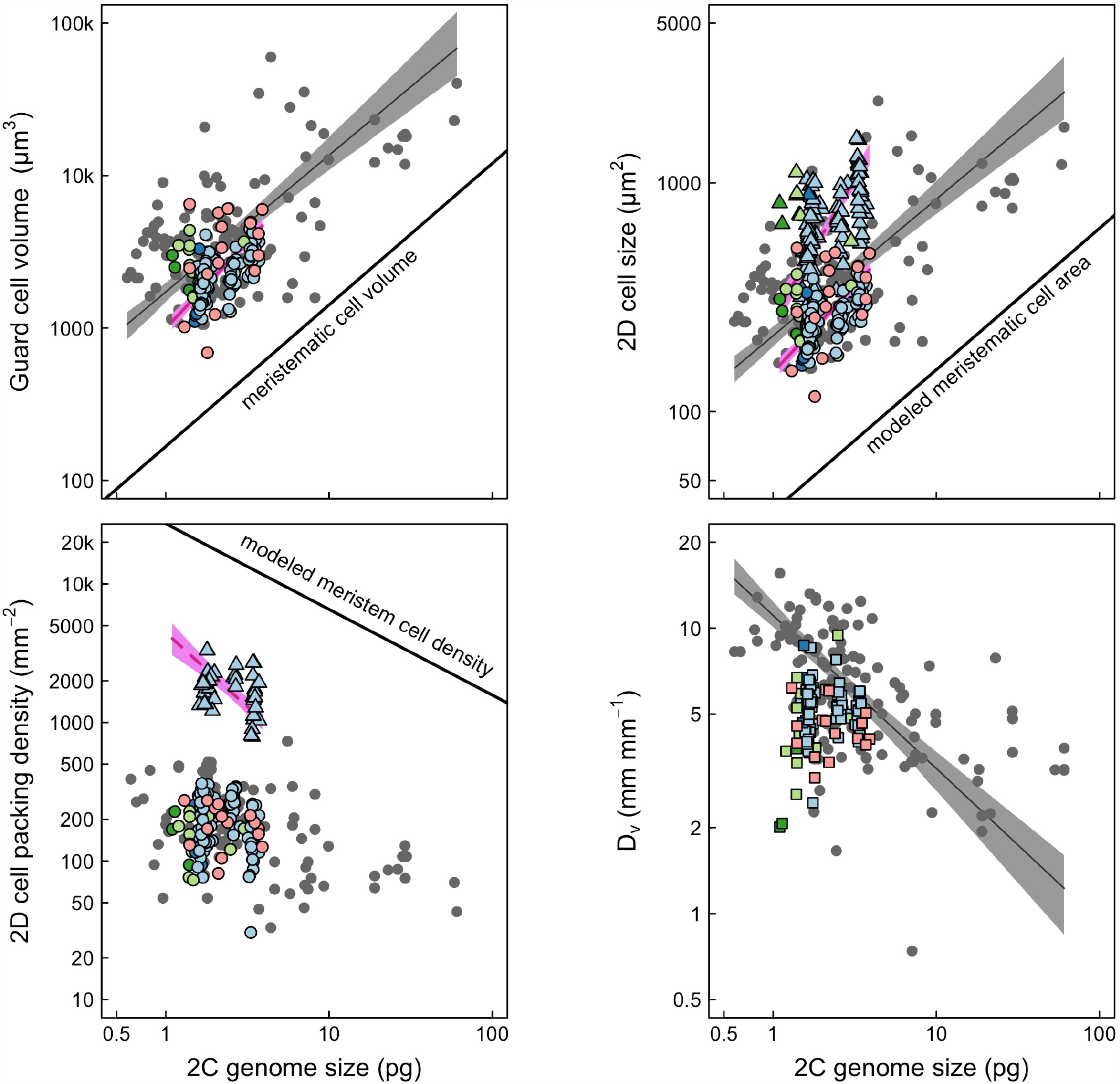
The effects of genome size on cell sizes and cell packing densities: (a) guard cell volume, (b) two-dimensional guard cell and epidermal cell sizes, (c) two-dimensional guard cell and epidermal cell packing densities, (d) leaf vein density. In all panels, the black solid line represents the meristematic cell volume, two-dimensional area, or two-dimensional cell packing density calculated from the scaling relationship between genome size and meristematic cell volume (Simova and Herben 2012) assuming spherical cells (see Theroux-Rancourt et al. 2021 for details). Grey points, lines, and shading represent the standard major axis regressions and 95% confidence intervals for the broad angiosperm dataset. Pink solid and dashed lines and shading represent standard major axis regressions and 95% confidence intervals of *Rhododendron* data. *Rhododendron* points are colored according to their subgenus: light blue = Pentanthera, dark blue = Ponticum, light green =-Rhododendron, dark green =-Tsutsusi, pink = Vireya. Point symbols indicate cell type: circles = guard cells, triangles = epidermal cells, squares = veins.

While vein density scaled negatively with genome size among angiosperms (*R*_*2*_ = 0.30, *P* < 0.0001), there was no effect of genome size on vein density among *Rhododendron* (*P* = 0.15, Figure 1d).

### Scaling among morphological traits

Leaf dry mass per area (*LMA*) can be driven by both variation in leaf thickness and leaf density. Among *Rhododendron*, lamina thickness (*L*_*T*_) scaled strongly with *LMA* (*R*_*2*_ = 0.76, *P* < 0.0001; Figure 2a). Leaf density (*LD*) was almost as strongly linked to *LMA* as lamina thickness (*R*_*2*_ = 0.70, *P* < 0.0001; Figure 1b). Leaf thickness and leaf density also scaled positively with each other (*R*_*2*_ = 0.23 *P* < 0.0001; Figure 2c). Because *LMA* can be influenced by both leaf thickness and leaf density, we partitioned the variance in *LMA* explained by these two variables. Among *Rhododendron*, leaf thickness explained 28% of the variation in *LMA*, and leaf density explained 19% of the variation in *LMA*. Jointly, leaf thickness and leaf density explained 50% of the variation in *LMA*, i.e. 50% of the variation in *LMA* cannot be separately attributed to either leaf density or leaf thickness.

**Figure 2.**
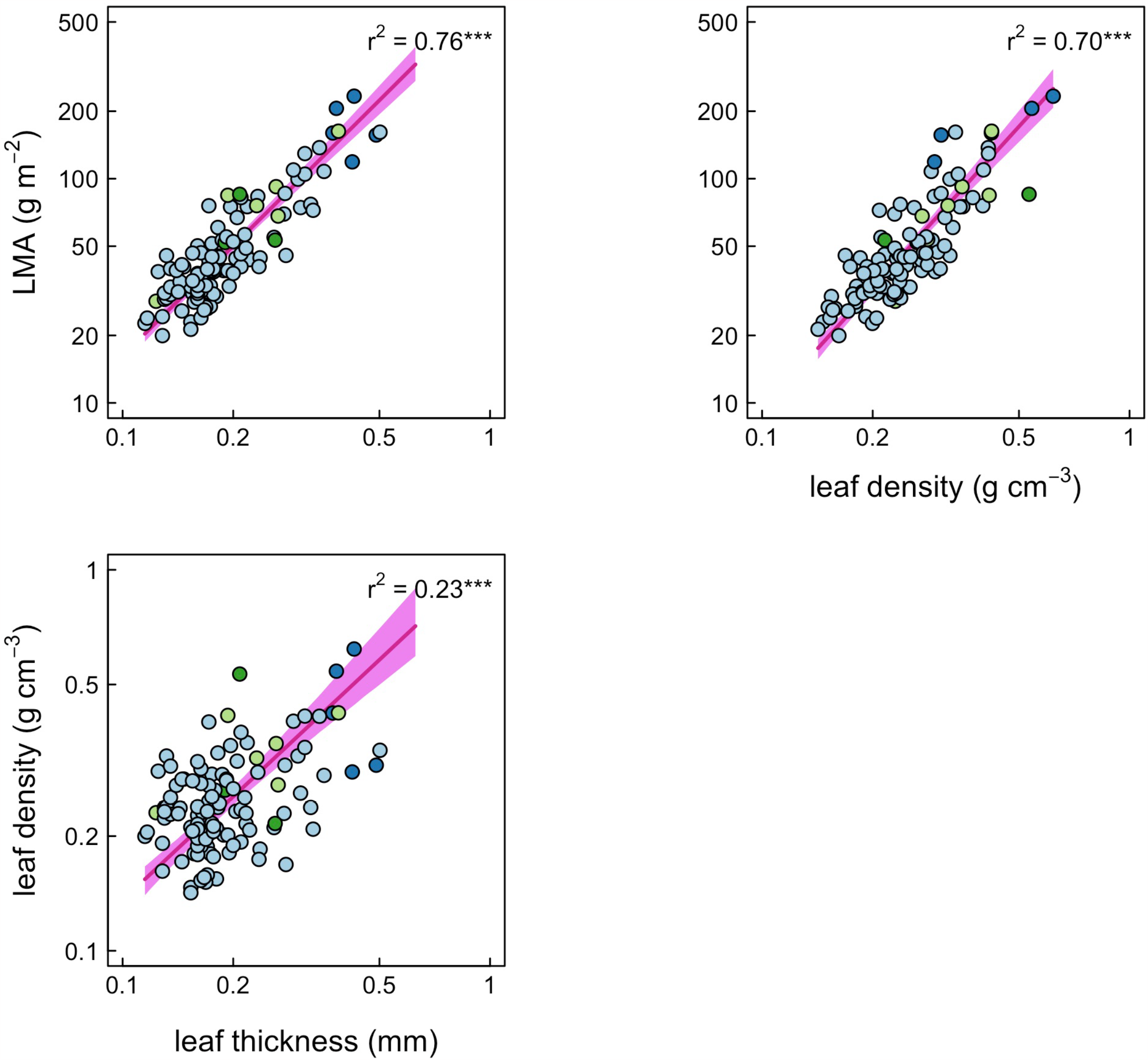
The scaling relationships between leaf thickness, leaf density, and leaf mass per area (LMA) for *Rhododendron*. Pink lines and shading represent standard major axis regressions and 95% confidence intervals. Points are colored according to clade (see Figure 1).

Because leaf water content has a strong effect on leaf function, we examined how leaf morphological traits scaled with *W*_*area*_ and *W*_*prop*_ (Figure 3). *W*_*area*_ scaled positively and significantly with lamina thickness (*R*_*2*_ = 0.86, *P* < 0.0001; Figure 3a), *LMA* (*R*_*2*_ = 0.69, *P* < 0.0001; Figure 3b), and leaf density (*R*_*2*_ = 0.23, *P* < 0.0001; Figure 3c). *W*_*prop*_, however, scaled negatively with increasing lamina thickness (*R*_*2*_ = 0.28, *P* < 0.0001; Figure 3d), *LMA* (*R*_*2*_ = 0.70, *P* < 0.0001; Figure 3d), and leaf density (*R*_*2*_ = 0.84, *P* < 0.0001; Figure 3d).

**Figure 3.**
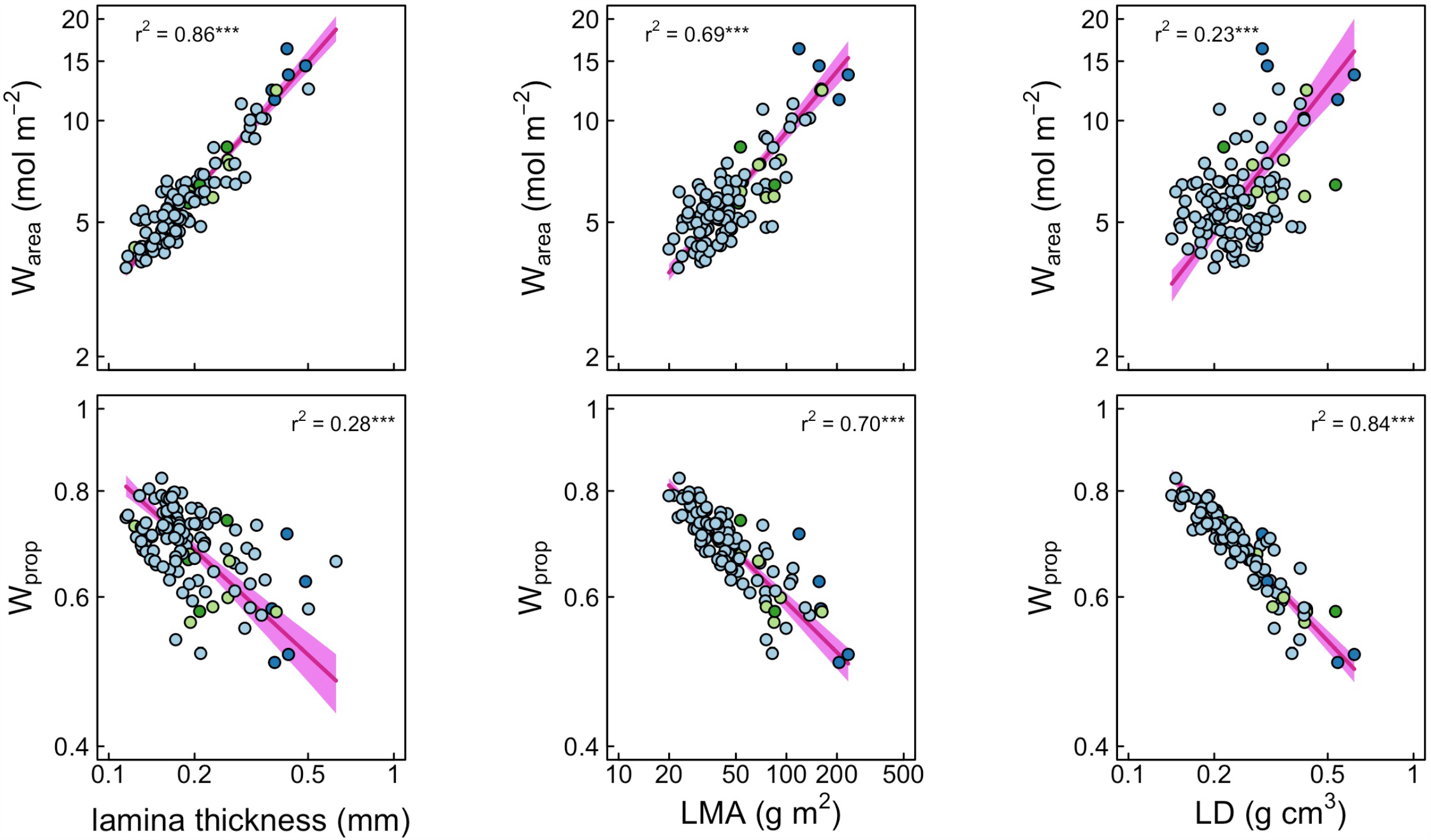
The effects of (a,d) leaf thickness, (b,e) *LMA*, and (c,f) leaf density (*LD*) on (a-c) water content per unit area (*W*_*area*_) and (d-f) water content per total leaf fresh mass (*W*_*prop*_). Pink lines and shading represent standard major axis regressions and 95% confidence intervals. Points are colored according to clade (see Figure 1).

### Effects of anatomical and morphological traits on leaf function

Leaf water residence time (τ) is a function of both stomatal conductance and water content, which is influenced by leaf morphology. There was a strong tradeoff between τ and *g*_*max,vol*_ (*R*_*2*_ = 0.95, *P* < 0.0001; Figure 4a), which was even stronger than the tradeoff between τ and *g*_*s,max*_ (*R*_*2*_ = 0.68, *P* < 0.0001; data not shown), most likely because normalizing by *g*_*s,max*_ by leaf volume incorporates *W*_*area*_, which is used in the calculation of τ. Both *W*_*prop*_ (*R*_*2*_ = 0.13, *P* < 0.001; Figure 4b) and *W*_*area*_ (*R*_*2*_ = 0.43, *P* < 0.0001; Figure 4c) also scaled with τ, as did lamina thickness (*R*_*2*_ = 0.33, *P* < 0.0001; Figure 4d) and *LMA* (*R*_*2*_ = 0.36, *P* < 0.0001; Figure 4e).

**Figure 4.**
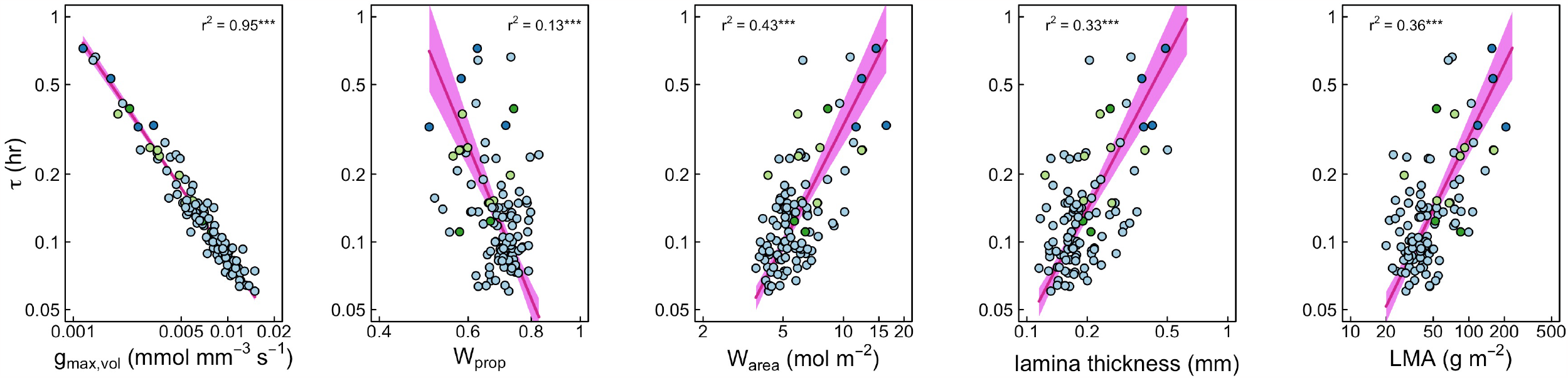
The responses of leaf water residence time (tau) to morphological and physiological traits: (a) modeled maximum stomatal conductance per leaf volume (*g*_*max,vol*_), (b) the water mass per leaf fresh mass (*W*_*prop*_), (c) leaf water content per unit leaf area (*W*_*area*_), (d) lamina thickness, (e) leaf mass per area (*LMA*). In all figures the pink line and shading represents the standard major axis regression and 95% confidence interval. Points are colored according to clade (see Figure 1).

**Figure 5.**
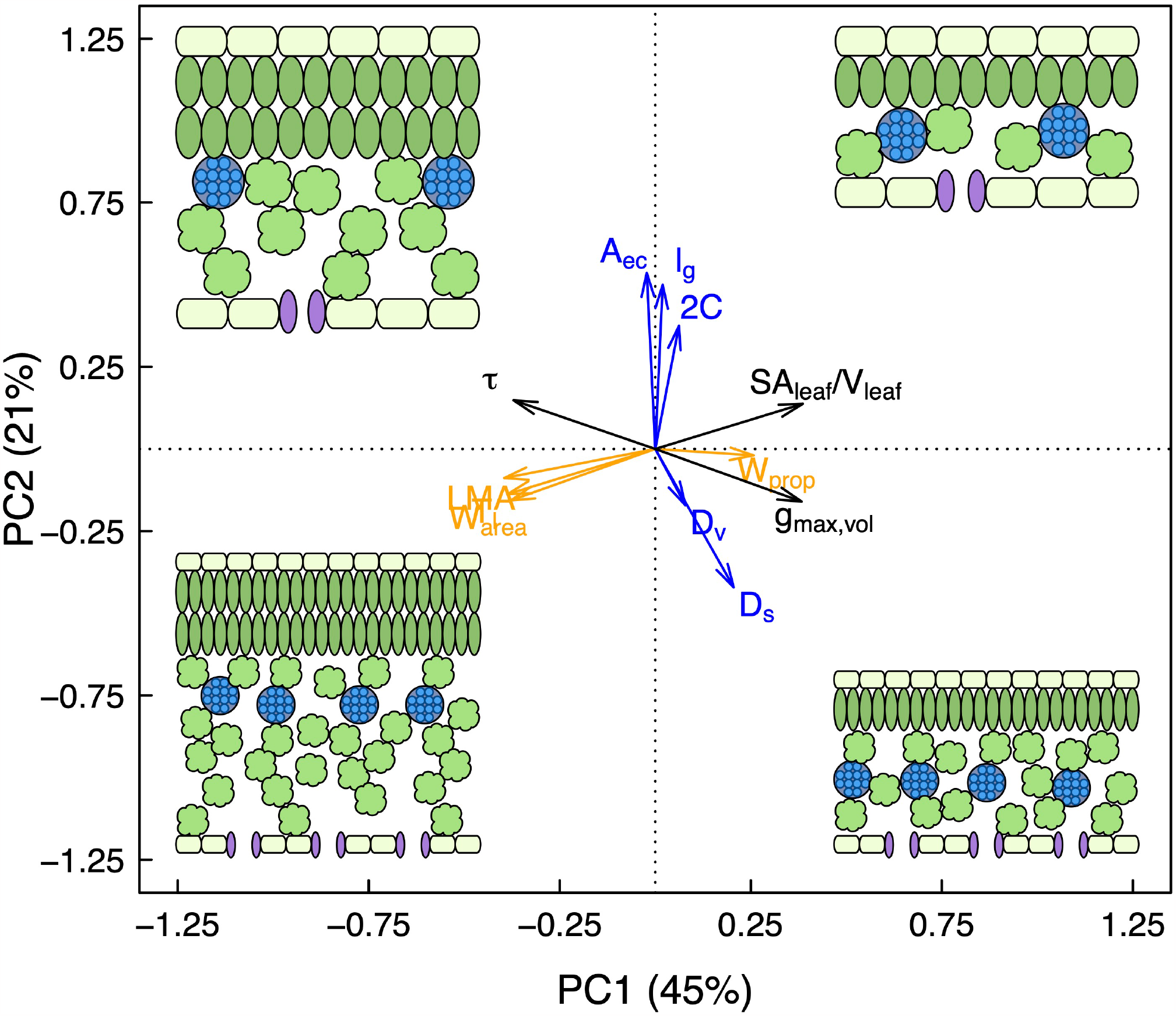
Principal components analysis of genome size and leaf traits among *Rhododendron* taxa. Variable loading vectors are colored according to trait type: blue are anatomical traits related to cell size and packing density, orange are morphological traits related to water content and dry mass investment, and black are physiological traits calculated from anatomical and morphological traits. Cartoon cross-sections of leaves in each quadrant illustrate the combinations of anatomical and morphological traits associated with leaves that would occur in that quadrant. Note that leaf thickness (*T*_*l*_) has almost identical loading to *LMA* and *W*_*area*_.

In multivariate space, anatomical traits related to genome size-cell size allometry and morphological traits related to leaf construction costs were largely orthogonal to each other (Figure 5). Almost half of the variation among *Rhododendron* leaves was driven by the first principal component, which was predominated by a tradeoff between high *W*_*prop*_ versus high *W*_*area*_, LMA and lamina thickness (Figures 2,3). Anatomical traits predominated the second principal component, which explained 21% of the variation among species, and was driven primarily by a tradeoff between large cells and genomes versus high packing densities (Figure 1). Because they are influenced by both anatomical and morphological traits, traits related to leaf function (*g*_*max,vol*_, τ, *SA*_*leaf*_*/V*_*leaf*_) were orthogonal to both morphological traits on PC1 and anatomical traits on PC2.

Because thinner leaves with higher *SA*_*leaf*_*/V*_*leaf*_ are capable of higher rates of energy and matter exchange than are thicker leaves, we tested how *SA*_*leaf*_*/V*_*leaf*_ scales with *g*_*max,vol*_ and τ and how these relationships are mediated by cell size. Among *Rhododendron, SA*_*leaf*_*/V*_*leaf*_ scaled positively with *g*_*max,vol*_ (*R*_*2*_ = 0.37, *P* < 0.0001; Figure 6a). The residuals of this relationship were negatively correlated with epidermal pavement cell size (*R*_*2*_ = 0.11, *P* < 0.0001; data not shown), implying that larger cells lead to lower *g*_*max,vol*_ for a given *SA*_*leaf*_*/V*_*leaf*_ (Figure 6b). Similarly, *SA*_*leaf*_*/V*_*leaf*_ scaled negatively with τ (*R*_*2*_ = 0.33, *P* < 0.0001; Figure 6c). The residuals of this relationship were significantly and positively correlated with epidermal pavement cell size (*R*_*2*_ = 0.08, *P* = 0.004; data not shown), implying that larger cells lead to longer τ for a given *SA*_*leaf*_*/V*_*leaf*_ (Figure 6d).

**Figure 6.**
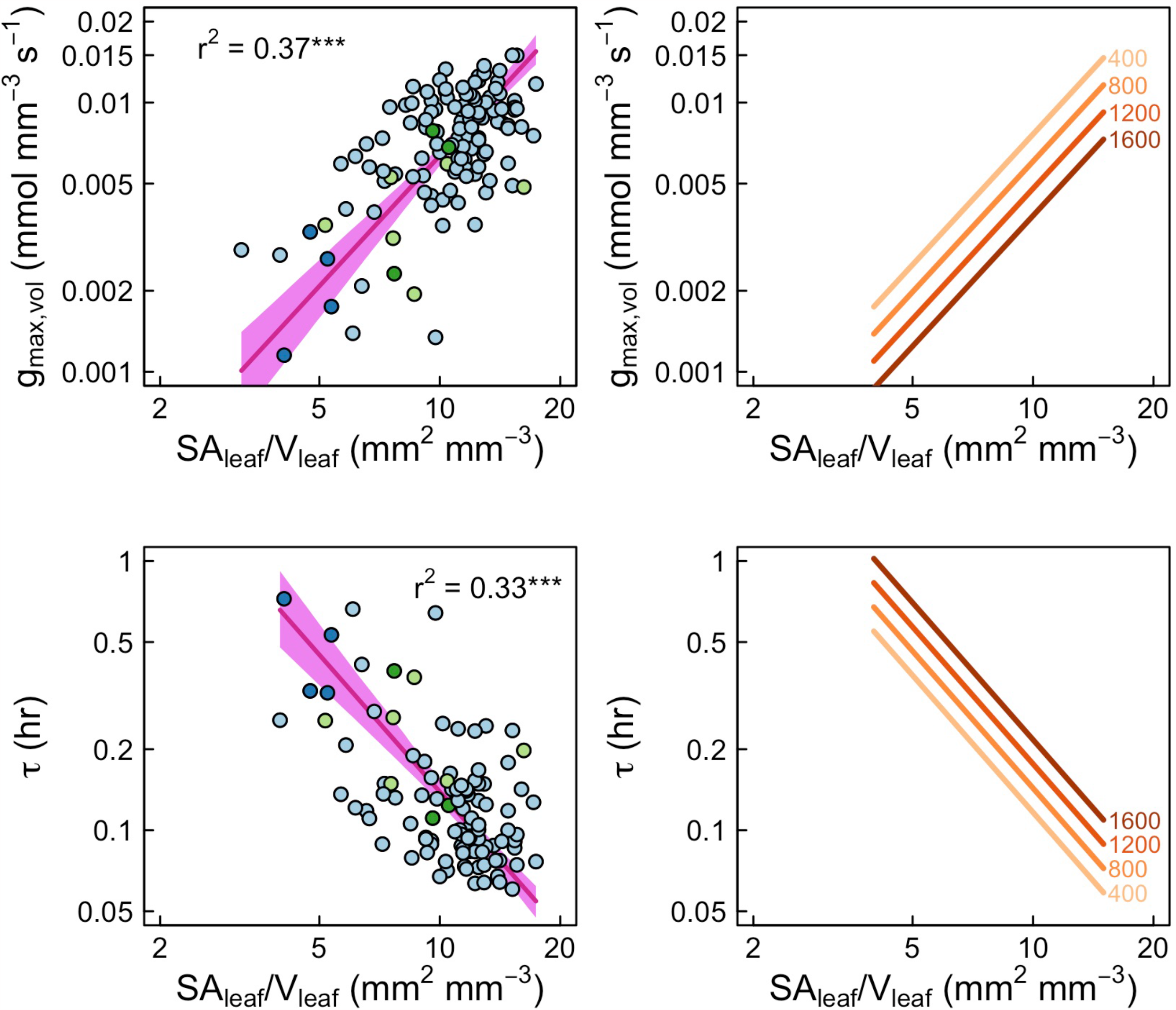
Both cell size and leaf morphology influence leaf functional traits. The effects of leaf surface area per volume (*SA*_*leaf*_*/V*_*leaf*_) on (a) maximum stomatal conductance per unit leaf volume (*g*_*max,vol*_) and (c) the leaf water residence time (τ). The residuals around these scaling relationships were significantly related to cell size. The effects of cell size on these residuals were used to model cell size isoclines (see Methods), which are shown in (b) and (d). Cell size (represented by 2D epidermal cell area in μm^2^) influences the scaling of *SA*_*leaf*_*/V*_*leaf*_ with (b) *g*_*max,vol*_ and (d) τ. In (a,c) the pink lines and shading represent the standard major axis regressions and 95% confidence intervals, with points colored according to their clade (see Figure 1). In (b,d) yellow-red lines indicate cell size isoclines with numbers adjacent to lines indicating the 2D epidermal cell sizes.

## Discussion

Using *Rhododendron* species as a case study, we show how genome size constraints on minimum cell size influence the construction and function of tissues and whole leaves. Our analysis clarifies how cell size variation influences whole leaf structure and function even though anatomical traits related to cell size and morphological traits related to construction costs can vary independently of each other. Furthermore, this analysis highlights how the maximum potential gas exchange of a leaf depends not only on the two-dimensional packing of stomata on the leaf surface, but more importantly on the three-dimensional structure of the leaf through which CO_2_ must diffuse. By using closely related congeneric species to illuminate these relationships, this study provides a powerful test of the effects of genome size on leaf structure without the confounding effects of large ecological and evolutionary differences often implicit in interspecific comparisons among phylogenetically diverse taxa.

### Effects of genome size on leaf anatomy

We found significant relationships between genome size, cell size, and cell packing density, consistent with previous analyses of phylogenetically broad sampling among angiosperms (Jeremy M Beaulieu *et al*. 2007b; Beaulieu *et al*. 2008; Simonin and Roddy 2018; Roddy *et al*. 2020; Théroux-Rancourt *et al*. 2021) and among diverse species within habitats (Jiang *et al*. 2023). Despite exhibiting relatively little interspecific variation in genome size compared to the total variation among angiosperms, *Rhododendron* species exhibited significant relationships between genome size, cell sizes, and cell packing densities (Figure 1). Overall, genome size was not as strong a predictor of packing densities as it was of cell sizes. Neither stomatal density (Figure 1b) nor leaf vein density (Figure 1d) were significantly related to genome size, in contrast to more phylogenetically diverse comparisons encompassing greater trait variation (Beaulieu *et al*. 2008; Simonin and Roddy 2018). That stomatal and epidermal cells were always larger and packed less densely than the limits imposed by meristematic cells is consistent with the fundamental effects of genome size on minimum cell size (Roddy *et al*. 2020). The presence of other cell types can also modify the effects of genome size on mature cell sizes and packing densities of any one tissue. For example, epidermal pavement cells fill the space unoccupied by stomata such that stomatal density is much lower than its potential maximum for a given stomatal size (Figure S2). Similarly, because multiple cell types occur in the leaf volume, there can be many combinations of cell sizes and packing densities among cell types (Roderick, Berry, Saunders, *et al*. 1999; John *et al*. 2013).

### Coordination among leaf morphological traits

There was strong coordination among morphological traits related to leaf construction. Because of the primacy of *LMA* in the LES, decomposing *LMA* into the factors driving it is important for understanding how leaves are built and function (Witkowski and Lamont 1991; Shipley 1995; Pyankov *et al*. 1999; John *et al*. 2017). Among *Rhododendron*, higher *LMA* was associated with both greater thickness and higher density, half of the variation in *LMA* was due to the joint effects of leaf density and leaf thickness that cannot be separated from each other (Figure 2).

Among *Rhododendron* leaf thickness explained a greater proportion of variation in *LMA* (28%) than did leaf density (19%), which is in contrast to previous analyses of Mediterranean woody species, among which 45% of the variation in *LMA* was due to leaf density and 33% to leaf thickness (De La Riva *et al*. 2016). Thus, increasing *LMA* in *Rhododendron* is due primarily to increasing thickness, either by larger cells typically associated with larger genome sizes or by additional layers of cells.

Because metabolism occurs in an aqueous environment and because water content influences hydraulic capacitance, higher water content links carbon economics and water relations [Roderick, Berry, Saunders, *et al*. (1999); Roddy *et al*. (2019); Huang *et al*. (2020); Wang *et al*. (2022); nadalIncorporatingPressureVolume2023]. All else being equal, thicker leaves have more leaf volume in which to have water, resulting in strong scaling relationships between leaf thickness, *LMA*, leaf density, *W*_*area*_, and *W*_*prop*_ (Figure 3), adding further support for the prediction that leaf morphology is associated with water content (Roderick, Berry, Saunders, *et al*. 1999). Though in global trait datasets photosynthetic rate and SLA plateau at higher leaf water contents (Huang *et al*. 2020; Wang *et al*. 2022), *LMA* and *W*_*prop*_ (equivalent to *LWC* in Huang et al., 2021) among *Rhododendron LMA* and *W*_*prop*_ are both in the range in which *SLA, W*_*prop*_, and photosynthetic rate exhibit strong positive scaling among global species. Thus, low *LMA* (high SLA) leaves have higher water content per dry mass that can support high *A*_*mass*_ (Wright *et al*. 2004).

In addition to its effects on leaf photosynthetic capacity, water content is also linked to carbon economics and leaf biomechanics. The combination of low *LMA* and high *W*_*prop*_ may indicate a greater reliance on a hydrostatic skeleton for biomechanical support, pattern that seems to extend also to flowers (Roddy *et al*. 2019; An *et al*. 2023; Roddy *et al*. 2023). Because the leaf mesophyll is under pressure from the leaf epidermis, the biomechanical properties of the living mesophyll tissue is critical to the maintenance of a porous mesophyll capable of high rates of CO_2_ diffusion (Théroux-Rancourt *et al*. 2021). In fact (and somewhat counterintuitively), this positive pressure imposed by the epidermis combined with the positive turgor pressure of the mesophyll is vital for the development of mesophyll porosity during leaf morphogenesis (Treado *et al*. 2022). Consistent with the role of mesophyll hydraulics in leaf biomechanics, leaves with low *LMA* shrink more during desiccation, suggesting that their high water contents are critical to their structural integrity (Scoffoni *et al*. 2014). High *W*_*prop*_ and low *LMA* may, therefore, facilitate higher photosynthetic rates and also cheaper leaves in terms of carbon, consistent with a strategy of fast growth and short leaf lifespan (Wright *et al*. 2004; Reich 2014; Roddy *et al*. 2023).

### Leaf construction and function from cells to whole leaves

Because there are many cell types in a leaf, variation at the level of the cell can impact higher order processes, such as whole leaf structure and photosynthetic capacity, though these effects can be diffuse. While genome size has a pervasive effect on all leaf cell types (Théroux-Rancourt *et al*. 2021), tissues can be modified independently of genome size-cell size allometry, either because cell sizes can be larger than the minimum bound imposed by genome size or because there are multiple cell types in the leaf (Figure 5). Individual cells can have thicker cell walls, increasing their mass at constant cell size, and cells can have various shapes that change the ratio of cell surface area to cell volume and ameliorate the limiting effects of large cell volumes on diffusion (Théroux-Rancourt *et al*. 2020; Treado *et al*. 2022). At the level of tissues, increasing thickness by adding more cell layers can influence tissue function independently of cell size and shape. Similarly, at the level of the leaf, leaf thickness and mesophyll porosity can vary independently of cell size and cell packing density allowing many leaf architectures to be built from the same cells (Figure 5) (Théroux-Rancourt *et al*. 2021). Leaf thickness (and, thus, *SA*_*leaf*_*/V*_*leaf*_), *LMA*, and *W*_*area*_ can be modified independently of cell size to influence photosynthetic capacity, leaf optics, and biomechanics. For example, sun leaves and shade leaves on the same plant that have the same genome size can vary dramatically in leaf thickness, *LMA*, and *W*_*area*_, resulting in different mesophyll structures for CO_2_ diffusion and different photosynthetic rates (Théroux-Rancourt *et al*. 2023). Our results highlight how morphological traits related to leaf construction vary orthogonally to anatomical traits related to cell size (Figure 5).

However, this does not mean that cell size has no effect on whole-leaf structure and function. Previous work has shown how smaller, more densely packed stomata in the epidermis allow for higher leaf surface conductance to CO_2_, which is critical to maintaining CO_2_ supply for photosynthesis during periods of declining atmospheric CO_2_ (Franks and Beerling 2009; Simonin and Roddy 2018). Once inside the leaf, CO_2_ must diffuse through the intercellular airspace and into the mesophyll cells. While smaller mesophyll cells allow for higher mesophyll surface area to be packed into a given leaf volume (Théroux-Rancourt *et al*. 2021), one of the major determinants of mesophyll conductance is leaf thickness, which directly impacts *SA*_*leaf*_*/V*_*leaf*_ (Roderick, Berry, Noble, *et al*. 1999; Earles *et al*. 2018). Though normalizing leaf surface conductance by leaf surface area is useful in characterizing the constraints on epidermal construction, area-normalized stomatal conductance does not account for the fact that each stoma must provide CO_2_ for an entire leaf volume, i.e. a stomatal vaporshed (Théroux-Rancourt *et al*. 2023). To account for the role of stomata in providing leaf volumes with CO_2_, we scaled maximum surface conductance by leaf volume (*g*_*max,vol*_). Because of the strong covariation among cell sizes throughout the leaf (Théroux-Rancourt *et al*. 2021), increases in *g*_*max,vol*_ due to reducing cell size would likely also be accompanied by increases in mesophyll conductance.

Because thinner leaves have a higher *SA*_*leaf*_*/V*_*leaf*_ available for energy exchange, we predicted that *SA*_*leaf*_*/V*_*leaf*_ would scale positively with *g*_*max,vol*_, indicating higher capacity for matter exchange with the atmosphere. Consistent with this prediction, there was a strong, positive relationship between *SA*_*leaf*_*/V*_*leaf*_ and *g*_*max,vol*_ (Figure 6a). Thus, as leaves become thinner and increase their *SA*_*leaf*_*/V*_*leaf*_, they also build their epidermis to better supply CO_2_ to their interior mesophyll tissue. Furthermore, variation in this bivariate relationship was mediated by cell size, such that for a given *SA*_*leaf*_*/V*_*leaf*_, reducing cell size enables higher *g*_*max,vol*_ (Figure 6b). Even though anatomical and morphological traits varied largely orthogonally to each other (Figure 5), cell size nonetheless impacts higher order leaf structure and function.

Leaves must optimize water loss and carbon gain under rapidly changing, dynamic environmental conditions. Though rarely measured, leaf water residence (τ) time is an important and dynamic indicator of how rapidly leaf physiology responds to changes in environmental conditions and reaches steady-state physiology, providing a functional link between water content and stomatal dynamics (Nobel and Jordan 1983; Hunt and Nobel 1987; Farquhar and Cernusak 2005; Simonin *et al*. 2013; Roddy *et al*. 2018). One of the strongest tradeoffs we observed among traits was the negative relationship between *g*_*max,vol*_ and τ, the leaf water residence time (Figure 4). The strength of the *g*_*max,vol*_-τ relationship is notable because τ exhibited a weaker relationship with *g*_*s,max*_, from which τ is calculated. We also predicted that *SA*_*leaf*_*/V*_*leaf*_ would be negatively related to τ because the greater capacity for energy exchange with the atmosphere associated with thinner leaves (higher *SA*_*leaf*_*/V*_*leaf*_) would allow for faster responses to the environment. Consistent with this prediction, *SA*_*leaf*_*/V*_*leaf*_ scaled negatively with τ (Figure 6c). Thus, thinner leaves have a higher capacity for energy exchange with the atmosphere (higher *SA*_*leaf*_*/V*_*leaf*_), have epidermises built to delivery higher fluxes of CO_2_ throughout their mesophyll tissue (higher *g*_*max,vol*_), and can more rapidly respond to fluctuations in environmental conditions (shorter τ). Furthermore, smaller cells allow for both higher *g*_*max,vol*_ and shorter τ (Figure 6c,d), thereby influencing whole leaf function. Even at constant leaf thickness, smaller cells would be predicted to be favored when high gas exchange rates and/or rapid responses to the environment are necessary.

The strong negative scaling between τ and *g*_*max,vol*_ (Figures 4, 5) suggests that these two variables more fully represent the fast-slow continuum of resource acquisition, resource use, and resource conservation in leaf traits proposed by Reich (2014) than traits typically measured as part of the LES. If being ‘fast’ at acquiring any particular resource to sustain growth (e.g. carbon, water, nutrients) requires being fast at processing all necessary resources (Reich 2014), then *g*_*max,vol*_ and τ can be used as indices of resource acquisition and conservation, respectively. While these two traits are driven predominantly by leaf morphology, they are nonetheless influenced by cell size (Figure 6), implying a role for cell size variation in determining ecophysiological strategies. The development of the LES has largely ignored anatomical traits related to cell size, though decomposing *LMA* into its constituent parts can include cell size and number (Poorter *et al*. 2009). Previous attempts to link genome size to the LES have revealed no significant relationships, despite the effects of genome size on cell size and maximum area-based photosynthetic rate (Wright *et al*. 2004; Jeremy M Beaulieu *et al*. 2007a; Simonin and Roddy 2018; Roddy *et al*. 2020). By determining the upper limit to leaf surface conductance (i.e. *g*_*s,max*_), genome size-cell size allometry defines the maximum limit of photosynthesis per unit leaf projected surface area (Roddy *et al*. 2020). Our analysis further clarifies how genome size-cell size allometry influences whole-leaf construction and the LES by helping to determine the maximum CO_2_ diffusion per unit leaf volume (*g*_*max,vol*_) and the speed of leaf physiological responses to the environment (τ) even when leaf thickness remains constant (Figure 6).

Our analyses also clarify how functional tradeoffs can result from recurrent motifs in anatomical and morphological traits. Though anatomical and morphological traits varied orthogonally to each other (Figure 5), they resulted in a strong tradeoff between *g*_*max,vol*_ and τ (Figure 4). The multivariate axis defined by the tradeoff between *g*_*max,vol*_ and τ is dominated by combinations of small cells and thin leaves (high *g*_*max,vol*_) versus large cells and thick leaves (high τ). This covariation in cell size and leaf thickness is consistent with theory and data showing that the vein density that optimally supplies leaf transpiration occurs when the distance between adjacent veins is equal to the distance between veins and the epidermis, i.e. vein packing densities should scale inversely with leaf thickness (Noblin *et al*. 2008). However, other analyses have shown that under different functional demands vein positioning may deviate from that predicted by optimal vein density (de Boer, Drake, *et al*. 2016). Our analyses for *Rhododendron* highlight that even though leaf thickness and vein density do not necessarily covary, other traits can interact with vein density and leaf thickness to drive an emergent functional tradeoff along the fast-slow continuum of leaf physiology. These analyses highlight how both whole-leaf and cell-level traits influence the maximum metabolic capacity of leaves and the rate of leaf responses to the environment.

Selection on different functional requirements can drive variation in traits orthogonal to the strong tradeoff between *g*_*max,vol*_ and τ. For example, thick leaves with high *LMA* are often tougher, providing better mechanical defense against leaf damage and better thermal capacitance, which may be beneficial in certain environments (Leigh *et al*. 2012; de Boer, Drake, *et al*. 2016; Tserej and Feeley 2021). These leaves may also be long-lived and tough, resistant to mechanical damage (Wright *et al*. 2004). Similarly, selection for increased CO_2_ supply to the mesophyll may result in smaller cells throughout the leaf even while leaf thickness and LMA remain constant because of selection for mechanical defenses. Yet, while cell size traits and leaf morphological traits can vary independently of each other (Figure 5), our analyses show that cell size nonetheless influences higher order leaf structure and function (Figure 6). In this way, cell size– and, indeed, genome size–may underlie these ecophysiological strategy axes, linking hydraulic, photosynthetic, and biomechanical functions, although there is substantial room for higher level modification of leaf construction to accommodate a range of genome sizes (Pyankov *et al*. 1999; Shipley *et al*. 2006; Roddy *et al*. 2020; Nadal *et al*. 2023). That these relationships exist among close relatives further reiterates the effects of genome size on leaf construction and ecological performance (Roddy *et al*. 2020).

## Acknowledgments

We thank Marc Hatchadourian and Claire Lyman of the New York Botanical Garden, Holly Forbes of the University of California Botanical Garden, and Steve Hootman of the Rhododendron Species Botanical Garden for facilitating access to and providing plant material. This work was supported by grants DEB-1838327 to K.A.S. and A.B.R. and CMMI-2029756 to A.B.R. from the U.S. National Science Foundation.

## Author Contributions

A.D., M.W., S.P., J.P., C.H., P.T., C.R., J.M., Y.-D. A., G.-F. J., and A.B.R. collected the data. A.D., K.A.S., and A.B.R. analyzed the data. A.B.R., A.D., and K.A.S. wrote the manuscript with input from all coauthors.

## Notes

### Competing Interest Statement

The authors have declared no competing interest.

