## Supplemental Information for "The effects of genome size on cell size and the functional composition and morphology of leaves: a case study in *Rhododendron* (Ericaceae)"

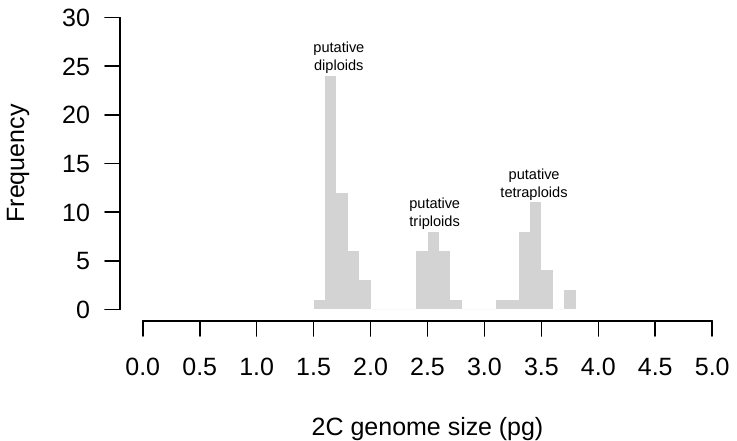

**Figure S1.** Histogram of 2C genome sizes for deciduous azalea samples, showing the three groups with putatively different ploidies.

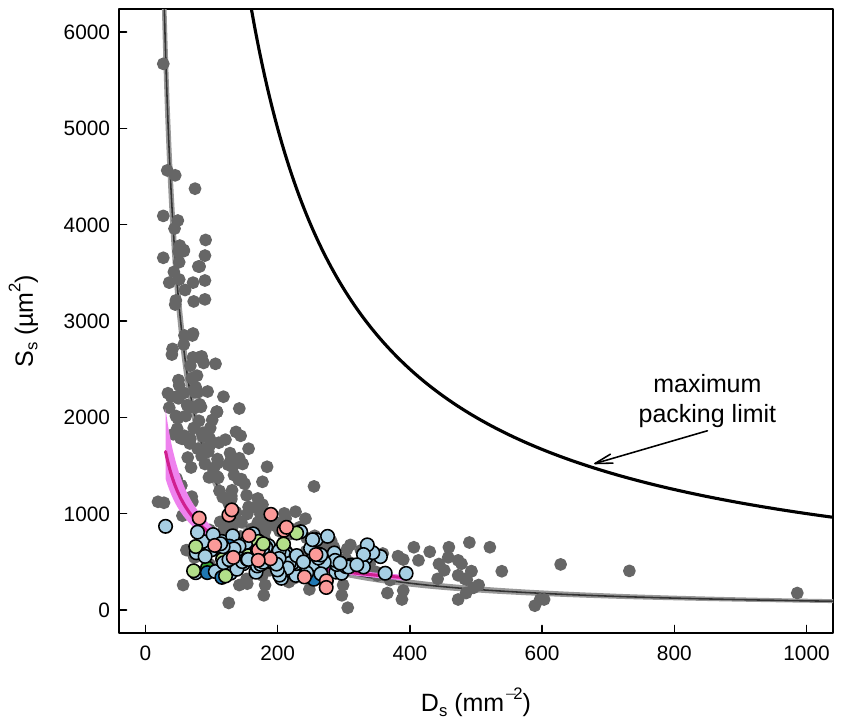

**Figure S2.** The tradeoff between stomatal size (Ss) and stomatal density (Ds). Grey points represent a broad sampling of angiosperms (see main text for details about data sources), and colored points represent *Rhododendron* leaves. The solid maximum packing limit line is where Ds = 1/Ss. Regression lines and shading represent standard major axis regressions (on log-transformed data) and 95% confidence intervals.

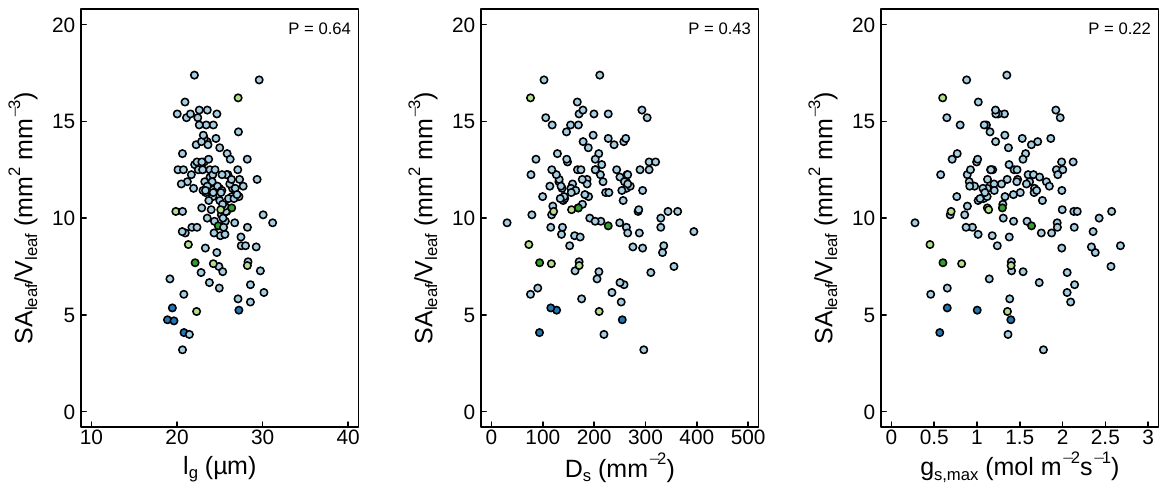

**Figure S3.** Relationships between guard cell length (*l_g_*), stomatal density (*D_s_*), and maximum stomatal conductance per unit area (*g_s,max_*) and leaf surface area to volume ratio (*SA_leaf_/V_leaf_* = 1/leaf thickness).

**Supplemental Information about ploidy and genome size among deciduous azaleas**

We sampled Rhododendron species known to vary in ploidy. These included the diploid species *alabamense*, *canescens*, and *viscosum* var. *ameulans*, the tetraploid species *austrinum* and *atlanticum*, three diploid elepidote species *arboreum* var. *arboreum*, *catawbiense* and *kyawii*, one diploid lepidote species *ciliicalyx*, and one diploid evergreen azalea species *pulchrum*. The breadth of sampling among deciduous azaleas was aimed at testing whether these species exhibit differences in ploidy (as determined by genome size differences) and hybridize to produce plants with intermediate genome sizes.

Based on previously published results and our preliminary data, we assumed ploidy from 2C genome sizes, with diploid azaleas ranging from 1.5-2.0 pg, triploids ranging from 2.4-2.8 pg, and tetraploids ranging from 3.1-3.8 pg (Figure S1). Previous studies have shown that evergreen species typically have genome sizes in the low end of the diploid range, elepidote species in the middle of the diploid range, and lepidote and deciduous azalea species in the middle to high end of the diploid range. The ranges of genome sizes for diploids, triploids, tetraploids, and pentaploids for the 65 deciduous azalea samples were also in line with those of previous studies (Supplementary Table 1). Two samples of hand crosses of seedlings from a diploid (seed parent) X tetraploid (pollen parent) had genome sizes in the tetraploid range. Two samples of hand crosses of seedlings from triploid (seed parent) X tetraploid (pollen parent) had genome sizes in the pentaploid range. One sample from a hand cross of pentaploid (seed parent) X tetraploid pollen parent had a genome size in the tetraploid range. All 17 naturally occurring hybrids had genome sizes in the triploid range and were found near diploid species (*R. canescens*, *R. alabamense*, and *R. viscosum* var. *ameulans*) and tetraploid *R. austrinum*.

Ploidy of deciduous azaleas has undergone recent revisions based on sampling of genome sizes using flow cytometry and chromosome counting. The deciduous azalea species *R. atlanticum*, *R. austrinum*, and *R. luteum* previously documented as diploid have been shown to be tetraploid, and *R. canadense* previously documented as tetraploid has been shown to be diploid. At the same time, what was previously identified as a late-blooming *R. alabamense* from the Red Hills region of Alabama and Georgia was found to be tetraploid, highlighting the characteristics that separated it from the diploid *R. alabamense*, resulting in it having a new species designation as *R. colemanii* (Zhou et al. 2008). Similarly, what was previously identified as the diploid *R. canescens* with very glandular new growth and pink flowers growing on the banks of the Escambia River, Blackwater River, and Yellow River near Pensacola, Florida, were found to be tetraploids, causing this group to be separated from the diploid *R. canescens* and moved into the tetraploid *R. austrinum* since this group had all the characteristics of *R. austrinum* with the exception of having pink flowers instead of yellow flowers (Miller 2011). Subsequent experimental crosses among azalea horticulturalists have shown that diploid (seed parent) X tetraploid (pollen parent) normally produce viable seed but tetraploid (seed parent) X diploid (pollen parent) fail to produce seedpods. Furthermore, flow cytometry measurements of seedlings from hand-crosses of diploid X tetraploid parents have shown that although most seedlings are triploid, some are tetraploid. These tetraploid seedlings as well as some of the triploid seedlings are fertile and produce seed when crossed with tetraploid pollen. Some fertile triploids produce seed when crossed with diploid or tetraploid pollen, and flow cytometry has shown seedlings from these crosses can range from diploid to pentaploid, including aneuploids between triploids and tetraploids. Flowering seedlings from diploid X tetraploid crosses have shown that most seedlings whether triploid or tetraploid have characteristics dominated by the tetraploid pollen parent including flower color.

Triploids in native interploidy contact zones between diploid deciduous azalea species and tetraploid *R. calendulaceum* have been well documented (Jones et al. 2007; Andrews 2014; Towe 2004; Li 1957). Preliminary results from flow cytometry suggested that triploids exist in interploidy contact zones of *R. calendulaceum* in South Carolina, North Carolina, Georgia, and West Virginia. The 17 samples in this study with genome sizes in the range of triploidy came from four interploidy contact zones for *R. austrinum* (one in Georgia and three in Florida). The contact zone in Albany, Georgia, contained the diploids *R. canescens*, *R. alabamense*, and *R. viscosum* var. *ameulans*. Two contact zones in Florida contained *R. canescens*, and one contact zone in Florida contained *R. alabamense*. The interploidy contact zones in Albany, GA, and Burnt Grocery Creek in Harold, FL, indicate that the triploids are found in closer proximity to diploid species than to tetraploid species and that these triploids have physical characteristics more similar to tetraploids, including in most cases flower color.

In general, field determination of ploidy at interploidy contact zones of deciduous azaleas based on visually identifiable traits (i.e. phenotypes intermediate between putative parental taxa) is often impossible. At contact zones between the diploid *R. canescens* and the tetraploid *R. austrinum*, all diploids have been pink-flowered and often eglandular on the new growth. Both tetraploids and plants subsequently identified as triploids are glandular on the new growth and can have yellow or pink flowers. Tetraploids are glandular on the new growth and can have yellow or pink flowers. Determining ploidy based on visual inspection at interploidy contact zones of the diploid *R. alabamense* and the tetraploid *R. colemanii* and at interploidy contact zones between the diploid *R. cumberlandense* and the tetraploid *R. calendulaceum* and between the diploid *R. viscosum* var. *ameulans* and the tetraploid *R. atlanticum* is even more challenging.
